# BINANA 2.0: Characterizing Protein/Ligand Interactions in Python and JavaScript

**DOI:** 10.1101/2021.09.10.459812

**Authors:** Jade Young, Neerja Garikipati, Jacob D. Durrant

## Abstract

BINding ANAlyzer (BINANA) is an algorithm for identifying and characterizing protein/ligand interactions and other factors that contribute to binding. We recently updated BINANA to make the algorithm more accessible to a broader audience. We have also ported the Python3 codebase to JavaScript, thus enabling BINANA analysis in the web browser. As proof of principle, we created a web-browser application so students and chemical-biology researchers can quickly visualize receptor/ligand complexes and their unique binding interactions.

## Introduction

Many biochemical processes depend on the association of specific proteins with their small-molecule ligands. The process by which a protein target recognizes its ligand (“molecular recognition”) is primarily determined by the non-covalent atomic interactions that form between the two, including hydrogen bonds, π-π stacking and cation-π interactions, electrostatic attraction and repulsion, and hydrophobics. These interactions contribute to the overall binding affinity of the protein/ligand association, and their geometric configuration plays a role in determining specificity (i.e., the tendency to bind the protein target but not other off-target proteins). Characterizing protein/ligand interactions can thus yield essential insights into the biological mechanisms underlying many processes (e.g., signaling, enzymatic catalysis, etc.). In the context of drug discovery, accurately characterizing protein/ligand interactions allows medicinal chemists to assess whether a ligand merits further study and pharmaceutical development.

When assessing a single protein/ligand complex, researchers often rely on manual inspection using visualization software such as VMD [1], PyMOL [2], or Chimera [3]. But many use cases require the assessment of many—sometimes thousands—of predicted ligand poses. The BINding ANAlyzer (BINANA) algorithm (first released in 2011) addresses this challenge by automating ligand-pose analysis [4], enabling the characterization of far more protein/ligand complexes than can be manually inspected. For example, McCarthy et al. [5] used BINANA to identify novel inhibitors of KRAS, a GTPase protein activated via mutation in 15% of human cancers. After performing a high-throughput virtual screen to evaluate six million compounds for potential KRAS inhibition, they used BINANA to identify the top predicted ligands that formed reasonable interactions with the protein receptor. These efforts ultimately led to an experimentally validated KRAS inhibitor. In a second study, Poli et al. used BINANA to identify inhibitors of monoacylglycerol lipase (MAGL) [6], a protein involved in the pathogenesis of neurodegenerative, cancer, inflammatory, and chronic-pain diseases. They docked ~14,000 molecules into the MAGL binding pocket and used BINANA to identify 17 compounds predicted to form critical interactions with the receptor. Subsequent experiments ultimately revealed three new compounds that inhibited MAGL activity; one even inhibited the proliferation of breast- and ovarian-cancer cell lines.

Several groups (including our own) have used BINANA to generate training data for machine-learning models (“scoring functions”) designed to identify ligands that merit more careful human scrutiny. Our NNScore2 algorithm [7] leverages BINANA descriptors (among other metrics) to predict ligand binding strength. NNScore2 has been used to help identify novel inhibitors of haloalkane dehalogenase [8], VEGFR-2 [9], and aromatase [10], among others. The DLSCORE scoring function [11] similarly uses BINANA descriptors to predict binding. BINANA has also been incorporated into several other programs (e.g., HBonanza [12] and POVME3 [13]), has inspired similar approaches [14, 15], and has been included in the Open Drug Discovery Toolkit [16].

These examples of broad adoption aside, the original BINANA implementation has some notable usability limitations. It runs only from the command line and provides no built-in visualization of the identified interactions, instead requiring separate visualization software. From the perspective of tool developers, BINANA 1.0 is also challenging because (1) its codebase organization does not allow for modular import into other Python scripts, (2) it is written in a now unmaintained programming language (Python2), and (3) its output is difficult to parse, complicating efforts to process BINANA analyses in other programs.

We developed BINANA 2.0 to address these challenges. The updated version can still run from the command line, but many users will benefit from our new web-browser implementation, which provides built-in molecular visualization that simplifies analysis. Tool-development researchers will benefit from updates to the Python codebase. We refactored the original implementation using a more modular programming approach that allows developers to integrate BINANA functions more easily into their projects (e.g., by importing individual modules as needed). We also added JSON-formatted output for easy processing by other computational tools and rewrote the code to be compatible with Python3 and JavaScript transpilation. To further encourage broad adoption and integration, we release BINANA 2.0 under a more permissive license than previous versions (Apache License, Version 2.0). Users can download the source free of charge from http://durrantlab.com/binana-download/ or access the browser app at http://durrantlab.com/binana/.

### BINANA Python codebase

#### Improving modularity and Python3 compatibility

We split the BINANA codebase into modules (separate files) to enable access as a Python library from other scripts. The original BINANA 1.0 was a stand-alone application (i.e., its codebase was contained in a single file, and its functions were not organized into modules), but several other groups have nevertheless incorporated BINANA code into their software projects [11, 13, 16]. To further enable such use, we refactored the BINANA codebase so other software can more easily import BINANA’s essential functions, including (1) loading PDBQT and PDB files containing receptor and bound-ligand structures, (2) analyzing those structures to identify specific protein/ligand interactions, and (3) saving BINANA analyses in various formats.

In refactoring the BINANA code, we also updated the codebase to make it compatible with Python3. The original version of BINANA was written in the now discontinued Python2 language.

#### Documentation

We created a documentation website to further improve BINANA usability: http://durrantlab.com/apps/binana/docs/. The website describes how to use the stand-alone BINANA program. It also catalogs the extensive docstrings associated with each public BINANA function so tool developers can quickly learn how to access the library’s application programming interface (API). Finally, the documentation also provides a copy of a Jupyter notebook demonstrating how to use BINANA as a Python library. The BINANA download includes the same notebook in an “examples” directory.

#### JSON output

The original version of BINANA saved binding-pose analyses to a PDB file or a VMD state file (for visualization using the popular program Visual Molecular Dynamics [1]). The new version of BINANA retains these features and further allows data export to the machine-readable JSON format. Many researchers have used BINANA to automatically assess the binding poses of large compound sets (e.g., in the context of virtual screens [5, 6]). To extract the data from these many analyses for subsequent processing, they have had to parse the BINANA-log text files directly. Now that BINANA outputs to JSON, this process will be much simplified.

#### Identifying protein/ligand interactions and other characterizations

To analyze a given protein/ligand complex, the user provides BINANA with molecular models of the protein and bound ligand in the PDBQT (recommended) or PDB format. BINANA then considers the positions and angles of various chemical groups to identify common interactions and otherwise characterize the complex.

BINANA 2.0 identifies the same protein/ligand interactions and characterizations that previous versions identified. These include close (< 4.0 Å by default) and closest (< 2.5 Å) contacts as well as hydrophobic, salt-bridge, π-π, and cation-π interactions. BINANA also tallies the number of times a ligand atom comes near the backbone or side chain of an alpha-helix, beta-sheet, or “other” secondary-structure amino acid. If the receptor/ligand models include hydrogen atoms, BINANA identifies hydrogen bonds. If the user provides models in the PDBQT format (which includes AutoDock atom types and Gasteiger partial charges [17]), BINANA also tallies the electrostatic energies between proximate protein/ligand atoms, the ligand atom types, and the number of ligand rotatable bonds. Full details can be found in the original BINANA manuscript [4].

The BINANA 2.0 interaction criteria are identical to the original version, except for close and closest contacts. Previously, these two interactions were mutually exclusive (i.e., those protein/ligand atom pairs that were close enough to be categorized as “closest” were not also considered to be “close”). In BINANA 2.0, all closest contacts are also close.

### BINANA JavaScript codebase and browser-app implementation

#### Porting BINANA to JavaScript

To broaden the impact of our BINANA algorithm, we transpiled the Python code to JavaScript using a software tool called Transcrypt (transcrypt.org). Transpilation rewrites or “translates” computer code written in one language (e.g., Python) into another (e.g., JavaScript). The resulting JavaScript library, BINANA.js, has the same functionality as the Python version but can be easily accessed from web apps running in any modern web browser. The BINANA download includes a Jupyter notebook and a simple HTML example file showing how to use BINANA.js.

#### BINANA browser app

To help non-computationalists better engage with the library, we integrated BINANA.js into a user-friendly browser-based application that detects and visualizes protein/ligand interactions.

##### Designing and compiling the browser-app user interface

The BINANA browser app provides an interactive graphical user interface (GUI). We designed the BINANA GUI using the same approach described elsewhere [18, 19]. In brief, the GUI was written in the TypeScript programming language, which compiles to JavaScript. We used Vue.js, an open-source web application framework, to compose reusable GUI components (e.g., text fields, buttons, etc.), and the BootstrapVue library to style all GUI components consistently according to the Bootstrap4 framework. We also used a custom molecular-visualization Vue.js component that leverages the 3Dmol.js JavaScript library [20] to display macromolecular and small-molecule structures in the browser, as required for visualizing BINANA-predicted protein/ligand interactions.

To compile these components and the BINANA.js library itself into a single web app, we used Webpack, an open-source module bundler, to manage the organization and composition of our source libraries and files. Webpack copies required files, combines files where possible, removes unneeded code, etc. The build process also used Google’s Closure Compiler to optimize the file size and performance of the TypeScript-compiled JavaScript code.

##### Browser-app Usage

###### Advanced parameters

The “Advanced Parameters” button appears at the top of the BINANA browser-app interface (Figure 1A). When clicked, a series of text fields appears that allows the user to modify the BINANA-library parameters. These fields initially contain the default values used by the BINANA command-line tool and Python library. We expect most users will wish to leave them unchanged, so they are hidden by default.

**Figure 1.**
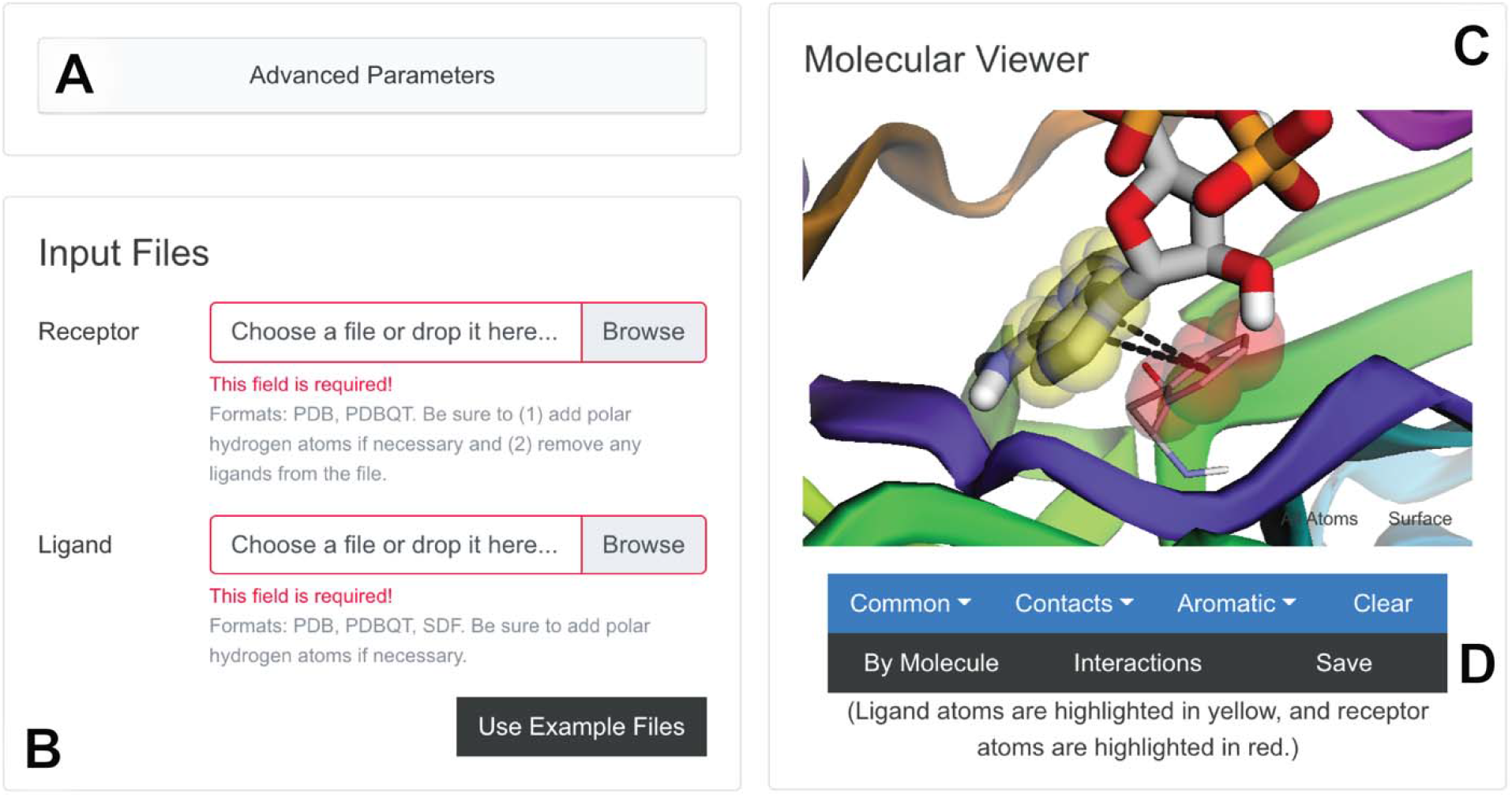
The BINANA web-app interface. (A) The “Advanced Parameters” button allows users to specify custom BINANA parameters. (B) The “Input Files” panel allows users to load their protein/ligand structures into the browser’s memory. (C) The “Molecular “Viewer” panel shows the detected interactions. (D) The “Save” button saves the results to the user’s disk.

###### Input files

The “Input Files” section allows users to load a ligand or receptor PDBQT or PDB file into their browser’s memory (Figure 1B). The structures are never uploaded to any third-party server, helping to ensure data privacy. Users can also run BINANA using build-in example ligand and receptor files by clicking the “Use Example Files” button.

###### Molecular viewer

The protein/ligand complex appears in the “Molecular Viewer” section of the browser app (Figure 1C), which also presents three general categories of interactions: “Common,” “Contacts,” and “Aromatic.” Clicking on the corresponding button opens a drop-down menu so users can choose which specific interaction to visualize. Under “Common,” users can select “Hydrogen Bonds,” “Hydrophobic,” or “Salt Bridge” interactions; under “Contacts,” users can select “Close” or “Closest” interactions; and under “Aromatic,” users can select “π-π Stacking,” “T Shaped,” or “Cation-π” interactions.

###### Interaction viewer

Once the user selects a specific interaction, semi-transparent red and yellow spheres highlight the participating protein and ligand atoms, respectively. We call this visualization scheme “By Molecule.” Clicking the corresponding button (Figure 1D) toggles on the alternate “By Interaction” scheme, in which hydrogen-bond donors and acceptors are highlighted in yellow and red, respectively, and salt-bridge positive and negative moieties are highlighted in blue and red, respectively. Clicking on the “Interactions” button further toggles the display of lines that connect the relevant protein and ligand atoms (Figure 1D). For hydrogen bonds, the line is a solid arrow that points from the hydrogen-bond donor to the acceptor. Otherwise, the line is dashed. The browser app also provides a “Save” button (Figure 1D) that allows users to save a zip file containing a copy of their protein/ligand files, as well as a thorough description of all BINANA.js-detected interactions in the JSON format, which is both human and machine readable.

## Examples of use

To test the web version of BINANA, we selected two receptor/ligand complexes and visualized them in the browser.

### M_2_ muscarinic acetylcholine receptor

Muscarinic acetylcholine receptors (mACHhRs) are G protein-coupled receptors (GPCRs) activated by acetylcholine [21]. As of 2017, ~34% of FDA-approved drugs targeted GPCRs [22], so studying GPCR/ligand complexes is useful for structure-based drug discovery and design [22]. Despite sharing between 64 and 82% sequence similarity, the five mACHhR subtypes differ in tissue distribution and GTP-binding protein partners [21]. The M_2_ muscarinic receptor, for example, is expressed in peripheral tissues and regulates heart rate [21]. Clinically approved small-molecule drugs that modulate M_2_ activity can effectively treat bradycardia (e.g., atropine [23]), urinary incontinence (e.g., tolterodine [23]), etc.

We used the BINANA web app to visualize a structure of M_2_ bound to LY2119620, a small-molecule, positive allosteric M_2_ modulator (Figure 2A). Allosteric muscarinic-receptor ligands are significant because they may enable improved receptor selectivity. All muscarinic receptors bind acetylcholine, so their orthosteric (primary) binding pockets are in many ways chemically similar. Identifying orthosteric ligands that bind to only one receptor subtype is thus challenging. In contrast, allosteric pockets may be more varied. Although LY2119620 also binds M_4_ muscarinic receptors and so is not strictly receptor specific [24], in principle allostery enables the design of ligands with improved specificity. We downloaded a PDB file of the receptor/ligand complex (PDB 6OIK [21]), added hydrogen atoms to the structure and ligand using MolProbity [25–27], and loaded the resulting model into the BINANA web app.

**Figure 2.**
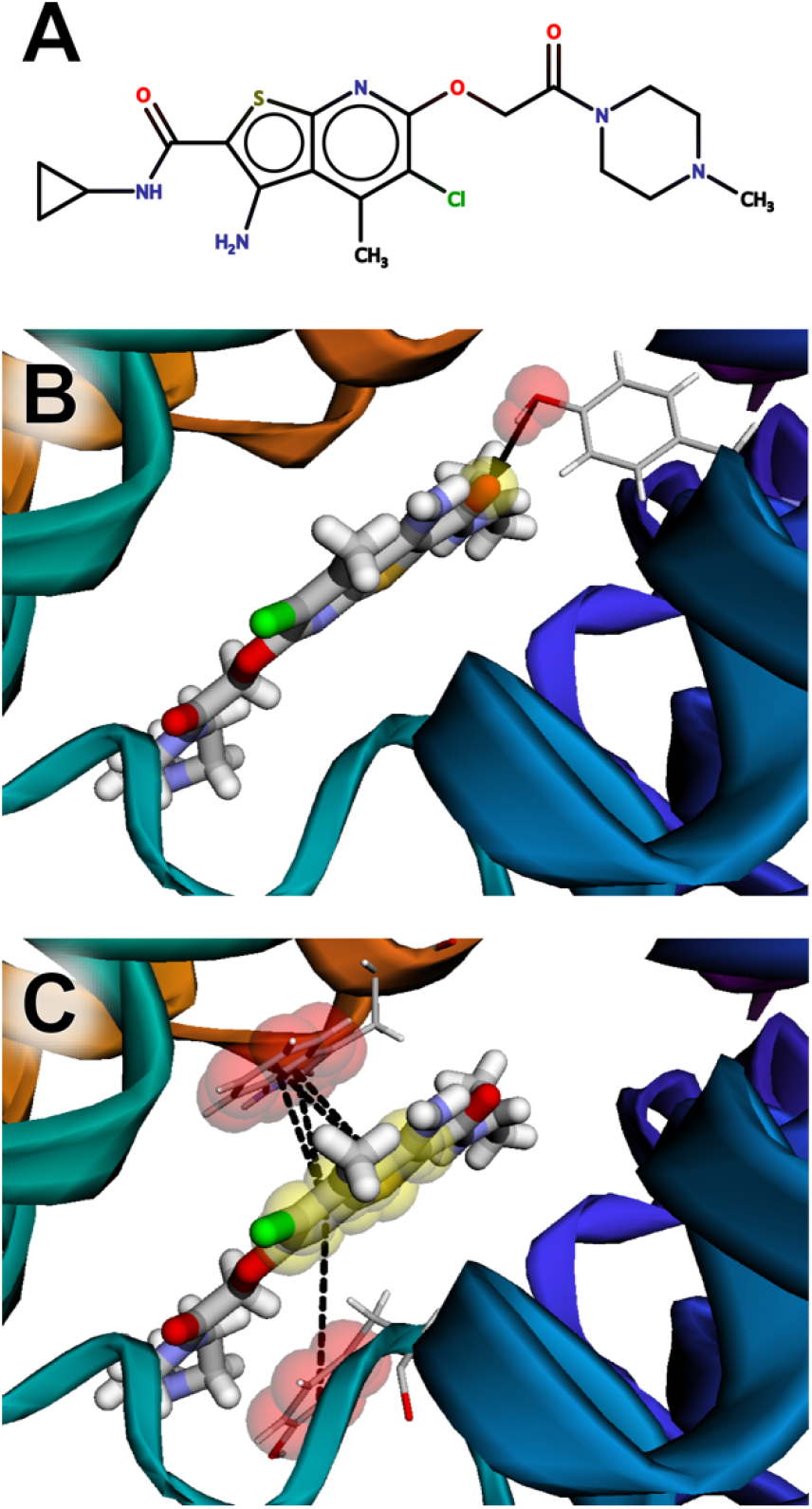
The muscarinic acetylcholine receptor M_2_ bound to LY2119620, an allosteric ligand. (A) A schematic of the ligand, created using MarvinSketch 18.24.0, ChemAxon (https://www.chemaxon.com). (B) The ligand forms a hydrogen bond with TYR80. (C) The ligand also participates in π-stacking interactions with TRP422 and TYR177.

BINANA visualization revealed a hydrogen bond between an LY2119620 carbonyl oxygen atom and the side-chain hydroxyl group of TYR80 (Figure 2B, black arrow). The aromatic LY2119620 bicyclic moiety also forms π-π stacking interactions with TRP422 and TYR177 (Figure 2C, black dotted lines).

### *Pseudomonas aeruginosa* peptidyl-tRNA hydrolase

Peptidyl tRNA hydrolase (Pth) is a potential drug target found in multiple species of bacteria, including *Escherichia coli*, *Mycobacterium tuberculosis*, *Mycobacterium smegmatis*, and *Pseudomonas aeruginosa*. The process of mRNA translation produces a peptidyl-tRNA intermediate, but ribosomes often release this intermediate when mRNA translation stalls. Pth separates peptidyl-tRNA into free tRNA and peptide by cleaving the ester bond between the C-terminus of the peptide and the 2’ or 3’ hydroxyl group at the 3’ end of the tRNA [28]. This cleavage frees the tRNA and peptide for reuse. In the absence of Pth activity, peptidyl-tRNAs cannot be recycled, ultimately resulting in bacterial death [28]. Small-molecule Pth inhibitors thus have potential as antibacterial therapeutics.

To demonstrate the BINANA web app applied to docked (predicted) ligand poses, we performed a virtual screen targeting the Pth active site. In brief, we prepared a model of the Pth receptor from *Pseudomonas aeruginosa* based on the 4QBK crystal structure [29]. We used PDB2PQR [30–32] to add hydrogens atoms to the protein, OpenBabel [33] to convert the PQR file to PDB, and MGLTools [34] to convert the PDB file to PDBQT. Because Pth binds peptide-based compounds, we also prepared a virtual library of approximately 60,000 easy-to-synthesize dipeptides provided by the Distributed Drug Discovery (D3) program [35–39]. We used Gypsum-DL to generate 3D models of these compounds and to enumerate alternate protonation, chiral, and tautomeric states [39]. The dipeptide files were also converted to the PDBQT format using OpenBabel and MGLTools.

We performed an initial docking run using Webina [18], a browser-app version of the docking program AutoDock Vina [40]. We used this initial run to determine appropriate coordinates and dimensions for the docking box and to confirm that our docking protocol could generally recapture the crystallographic pose of a known ligand (PDB 4QBK [29]). Having identified acceptable parameters, we docked all ~60,000 compounds using command-line Vina running on resources provided by the University of Pittsburgh’s Center for Research Computing (default parameters).

The two best-scoring compounds both had Vina scores of −9.0 kcal/mol. We loaded one of these, (2S)-2-[(2R)-2-amino-3- (anthracen-9-yl)propanamido]-3-(quinolin-2-yl)propanoic acid, into the BINANA web app (Figure 3A). The visualization suggests that the ligand participates in three hydrogen bonds with the receptor, each with ASN116 (Figure 3B, black arrows), as well as a π-stacking interaction with TYR68 (Figure 3C, dotted lines).

**Figure 3.**
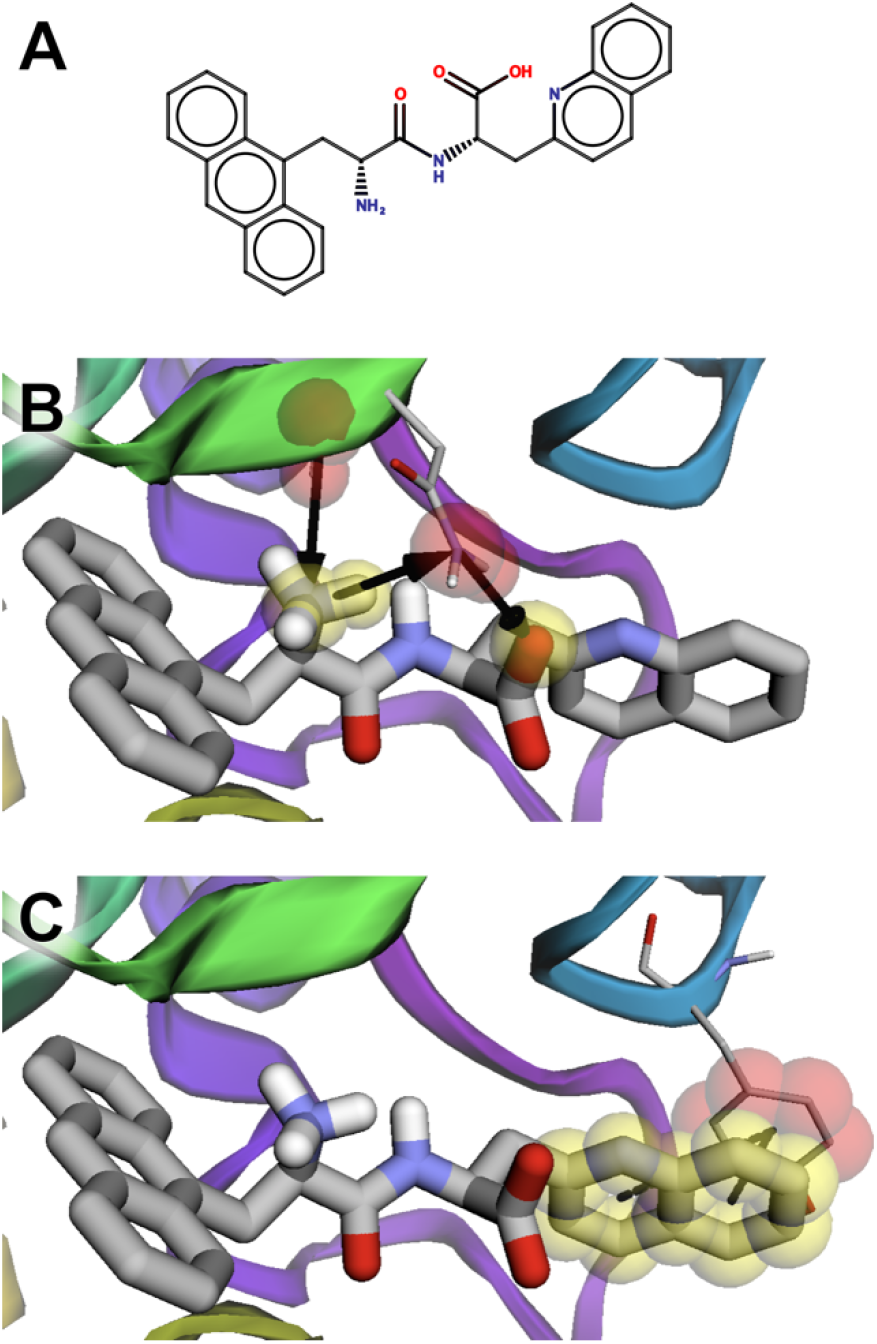
The peptidyl-tRNA hydrolase bound to a predicted ligand identified in a virtual screen. (A) A schematic of the ligand, created using MarvinSketch 18.24.0, ChemAxon (https://www.chemaxon.com). B) The ligand is predicted to form multiple hydrogen bonds with ASN116. (C) The ligand also participates in a π-stacking interaction with TYR68.

## Related programs

Several free desktop tools can also characterize receptor/ligand complexes (e.g., Visual Molecular Dynamics, PyMOL, and UCSF Chimera). Similar commercial tools include MOE (chemcomp.com), Discovery Studio (accelrys.com), SAMSON (samson-connect.net), and Small Molecule Drug Discovery Suite (schrodinger.com). Though powerful, these tools require separate download and installation. Some are also expensive, and even the free programs impose restrictions on commercial use in some cases. Furthermore, none of these desktop programs provides a JavaScript API that enables easy integration into user-friendly browser apps. In contrast, BINANA 2.0 includes a JavaScript implementation called BINANA.js, which we used to build a web app that can be freely accessed by simply visiting a website. We also release BINANA 2.0 under the terms of the open-source Apache License, Version 2.0, which permits incorporation into any program, commercial or otherwise.

nAPOLI [41] and PLIP [42] are examples of free online tools for characterizing receptor/ligand complexes. Both include convenient web-based interfaces, and PLIP also works as a command-line program. BINANA.js-powered browser apps have several advantages over these useful server apps. For example, BINANA.js allows apps to detect intermolecular interactions in the browser itself, without requiring users to upload their (possibly proprietary) structures to a third-party system. Instead, a simple web server sends the BINANA.js library to users’ browsers to detect interactions locally on their own machines. Consequently, BINANA.js-powered browser apps do not require an extensive remote computing infrastructure where calculations take place “in the cloud.”

## Broad compatibility

We have tested the BINANA Python and JavaScript libraries on the operating systems, Python versions, and browsers listed in Table 1. The software depends on no external Python or JavaScript libraries, so we do not anticipate compatibility issues on other, untested setups.

**Table 1.** Operating system, Python, and web-browser compatibility.

| Operating System | Python | Browser (JavaScript) |
| --- | --- | --- |
| Ubuntu (Linux) 20.04.2 LTS | Python 3.9.1 | Chromium 92.0.4515.159 |
|  |  | Firefox 91.0 |
| macOS Mojave 10.14.6 | Python 3.6.7 | Chrome 93.0.4577.42 |
|  |  | Firefox 91.0.1 |
|  |  | Safari 14.1.2 |
| Microsoft Windows 10 Home | Python 3.8.8 | Chrome 92.0.4515.159 |
|  |  | Firefox 91.0.1 |
|  |  | Edge 92.0.902.73 |
| Android 11 | N/A | Chrome 92.0.4515.131 |
|  |  | Firefox 90.1.1 |
| iOS 14.7.1 | N/A | Safari 14.1 |

## Conclusion

BINANA 2.0 retains the core functionality of the original version in that it can run as a stand-alone, command-line program. But it now also serves as a Python library that others can incorporate into their Python-based computational-biology tools. We also ported the BINANA library to JavaScript, enabling use in the web browser. To demonstrate how to incorporate BINANA.js into browser-based applications, we created the BINANA browser app. This app can be accessed online, enabling easy access and visualization without requiring command-line use.

## Data software and availability

Users can download the BINANA 2.0 source code—including the Python3/JavaScript libraries and the web-app graphical user interface—free of charge from http://durrantlab.com/binana-download/. We release BINANA 2.0 under the terms of the open-source Apache License, Version 2.0. Users can also freely access the BINANA browser app at http://durrantlab.com/binana/, and the API documentation at http://durrantlab.com/apps/binana/docs/.

## Supporting information

The “SMILES.csv” file includes the names and SMILES strings of the ligands depicted in Figures 2 and 3.

## Acknowledgements

We thank the University of Pittsburgh’s Center for Research Computing for providing helpful computer resources. This work was supported by the National Institute of General Medical Sciences of the National Institutes of Health [R01GM132353 to J. D. D.]. The content is solely the responsibility of the authors and does not necessarily represent the official views of the National Institutes of Health.

## Table of contents graphic

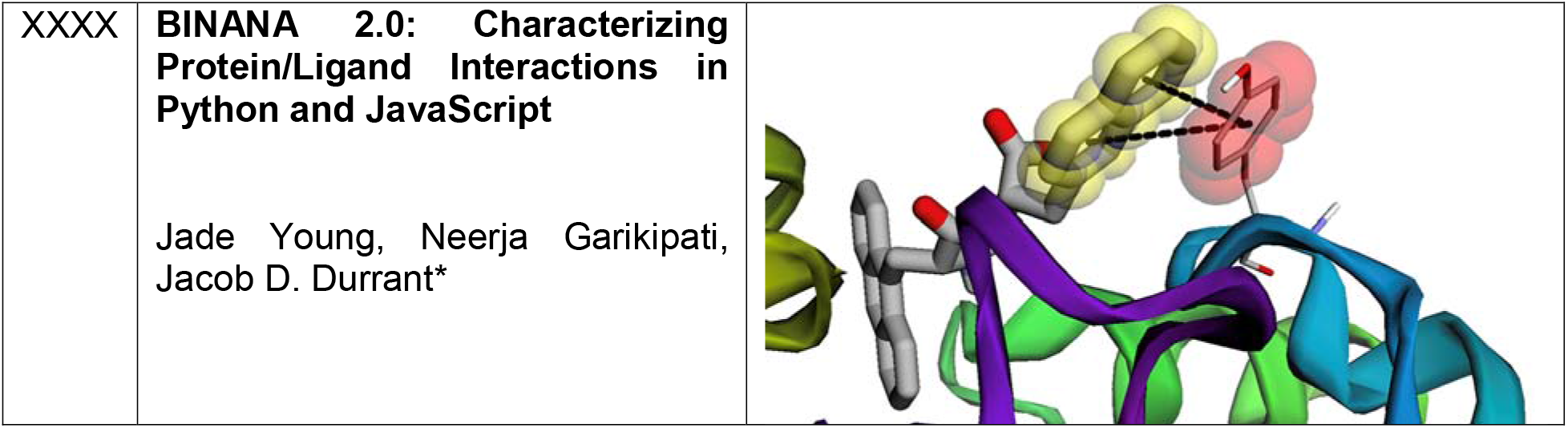

